# The genomic basis of hybrid male sterility in *Ficedula* flycatchers

**DOI:** 10.1101/2022.09.19.508503

**Authors:** J. Carolina Segami, Carina F Mugal, Catarina Cunha, Claudia Bergin, Monika Schmitz, Marie Semon, Anna Qvarnström

## Abstract

Identifying genes involved in genetic incompatibilities causing hybrid sterility or inviability is a long-standing challenge in speciation research, especially in studies based on natural hybrid zones. Here we present the first high-probability candidate genes for hybrid male sterility in birds by using a combination of whole genome sequence data, histology sections of testis and single cell transcriptomics of testis samples from male pied-, collared-, and hybrid flycatchers. We reveal failure of meiosis in hybrid males and propose candidate genes involved in genetic incompatibilities causing this failure. Based on identification of genes with non-synonymous fixed differences between the two species and revealing miss-expression patterns of these genes across the various stages of hybrid male spermatogenesis we conclude aberrant chromosome segregation and/or faulty chromatin packing. A lower proportion of spermatids produced by hybrid males implies that a proportion of the aberrant spermatids undergo apoptosis. Finally, we report an overrepresentation of Z-linkage of the revealed candidate incompatibility genes. Our results challenge the assumption that speciation processes are driven by fast evolving genes by showing that a few changes in genes with highly conserved and central functions may quickly ensure reproductive isolation through post-zygotic isolation.

## Introduction

In sexually reproducing organisms, evolutionary divergence that causes reproductive isolation is necessary to achieve speciation. Although there are several types of reproductive barriers that can generate and maintain genetic integrity of emerging new species, post zygotic barriers are considered to be more permanent, especially intrinsic postzygotic isolation in the form of hybrid sterility or inviability (Muller, 1942; Coyne & Orr, 2004; Coughlan & Matute, 2020). However, even if there is an extensive theoretical framework outlining possible explanations for how intrinsic postzygotic can arise, these ideas remain largely empirically untested. The molecular basis for specific genomic clashes still remains poorly known, with the exception of a few model systems. We therefore need to reveal the underlying genetics in a broader range of organisms if we want to understand the evolutionary forces driving speciation.

The well-established Bateson-Dobzhansky-Muller model (Bateson, 1909; Dobzhansky, 1936; Muller, 1942) proposes that hybrid intrinsic dysfunction is caused by negative epistatic interactions between two or more loci (i.e. BDMI’s). These loci have become incompatible with each other following divergence in different allopatric populations and cause a lethal or underperforming hybrid phenotype when these populations interbreed at secondary contact.

The common observation of greater intrinsic fitness reduction of hybrids belonging to the heterogametic sex, known as Haldane’s rule, implies a particular role of sex-linked genes in causing genetic incompatibilities (Coyne & Orr, 2004). The main hypotheses outlined to explain Haldane’s rule suggests a higher exposure of recessive incompatible alleles when hemizygous (Muller, 1940). This idea holds even when incompatibility genes are randomly distributed across the genome. Alternatively, there may be a relatively faster accumulation of incompatibilities on the sex-chromosomes as compared to the autosomes due to an overall relatively faster evolutionary divergence of sex-linked genes “faster-X”, (Charlesworth et al., 1987). In addition, genes that are especially crucial for functional reproduction may be located on the sex-chromosomes including potentially troublemaking genes such as selfish meiotic-drive alleles that can escape the control of the co-evolved suppressors in hybrids (Pomiankowski & Hurst, 1993). Finally, some additional ideas have been proposed that do not focus on sex chromosomes and cannot explain Haldane’s rule in species where females constitute the heterogametic sex. One such idea is that male reproductive traits in general may evolve faster than female reproductive traits due to more intense reproductive competition among males (Wu & Davis, 1993; Wu et al., 1996).

Few examples of Bateson-Dobzhansky-Muller incompatibilities (BDMIs) have been identified and most of them have been established by empirical work using model organisms such as drosophila and mouse (Presgraves, 2010; Maheshwari & Barbash, 2011). Based on these studies there is strong support for a non-random expression of intrinsic isolation on the X chromosome, but few studies control for recessively of hybrid dysfunction reviewed by (Qvarnström & Bailey, 2009; Charlesworth et al., 2018). However, the majority of the studies that have detected BDMIs have focused on species pairs than have diverged 2 or 3 million years ago and therefore may not naturally hybridize any more (Dobzhansky, 1936; Sawamura, 2000; Mallet, 2006; Presgraves & Meiklejohn, 2021). In order to obtain a comprehensive understanding on the role of BDMIs in speciation it is important to reveal the molecular basis to hybrid dysfunction in natural populations of closely related species that hybridize. Only two genes driving hybrid incompatibilities have been identified in vertebrates, *prdm9* for mouse (Mihola et al., 2009) causing hybrid sterility and *xmrk* in sword tail fish causing hybrid inviability (Gordon, 1937; Powell et al., 2020). There are, to our knowledge, no previous identified candidate genes for hybrid incompatibilities in birds where males constitute the homogametic sex. Male gametogenesis is a complicated process that involves many stages of development that could potentially fail in hybrids. Since this process also is encoded by a large percentage of the genome, the number of potential diverged genomic sites that may cause hybrid male sterility is high. Identifying high-probability BMDI candidates for hybrid male sterility therefore requires establishing high-resolution genotype-phenotype connections.

In this study, we use naturally hybridizing *Ficedula* flycatchers to shed novel light on the evolution of hybrid male sterility in a study system with well-developed genomic tools (Qvarnström et al., 2010, 2016). Collared and pied flycatchers are closely related species that diverged less than ~1 million years ago (Ellegren et al., 2012). The males differ in important sexually selected traits such as plumage coloration and song vocalization but mixed-species pairing regularly occur at natural hybrid zones. The most recently formed hybrid zone is found on the Baltic Island Öland where collared flycatchers colonized the island during the 1960s. Hybrids between collared and pied flycatchers display several intermediate phenotypic including plumage (Alatalo et al., 1982; Svedin et al., 2008) and song (Gelter, 1987; Qvarnström et al., 2010). Hybrid males also show clear signs of intrinsic incompatibilities in terms of higher metabolic rate (McFarlane et al., 2016), and malformed sperm (Alund et al., 2013). Mugal et al., 2020 found an intermediate gene expression of testis genes in bulk samples from hybrids indicating no major incompatibilities between collared and pied flycatchers. However, hybrids may have a different testis cell composition than the pure species (Hunnicutt et al., 2022) and we therefore recently characterized the spermatogenesis of *Ficedula* flycatchers at a single cell level to reveal species differences in gene expression at specific stages of the process (Segami et al., 2022). We found that most differentially expressed (DE) genes were active after meiosis and typically coding for cell respiratory or motility pathways at a late stage of spermatogenesis. However, we also detected DE genes during meiosis. Among these DE genes there was a tendency for enrichment of Z-linked genes suggesting fast Z evolution.

Here, we aim at identifying the molecular basis of sterility in *Ficedula* male hybrids by characterizing the hybrid spermatogenesis transcriptome at a single cell level. Using single-cell RNA sequencing of testis cell suspensions of two hybrids, three Collareds and three Pied flycatchers and markers developed by (Segami et al., 2022) and histology sections we first aim to investigate whether hybrid males go through all “normal” stages of spermatogenesis by identifying if hybrids have all testis cell types found in the parental species. Secondly, we will investigate whether hybrid expression patterns are different from the parental species in any particular stage of spermatogenesis as an indication of a possible failure. Finally, we identified non-synonymous fixed differences between the two species and analyzed their role across the various stages of spermatogenesis and connect this information with possible miss-expression patterns.

## Results

We generated scRNA data from testis cells suspensions obtained from 8 sacrificed male birds (3 Pied flycatchers, 3 collared flycatchers and 2 F1 hybrids, both with a pied mother, see Table S1). We created a consensus clustering using the 6 pure species samples to use as a baseline for all our consecutive analysis. The identities of the obtained 17 different cell clusters (Figure 1A) were verified using the characterized flycatcher markers for spermatogenesis populations developed by Segami et al 2022 (Figure S1). Our new clusters corresponded to all the previously identified testis cell clusters in flycatcher spermatogenesis except for one of the somatic cell clusters that was not found (Segami et al 2022).

**Figure 1.**
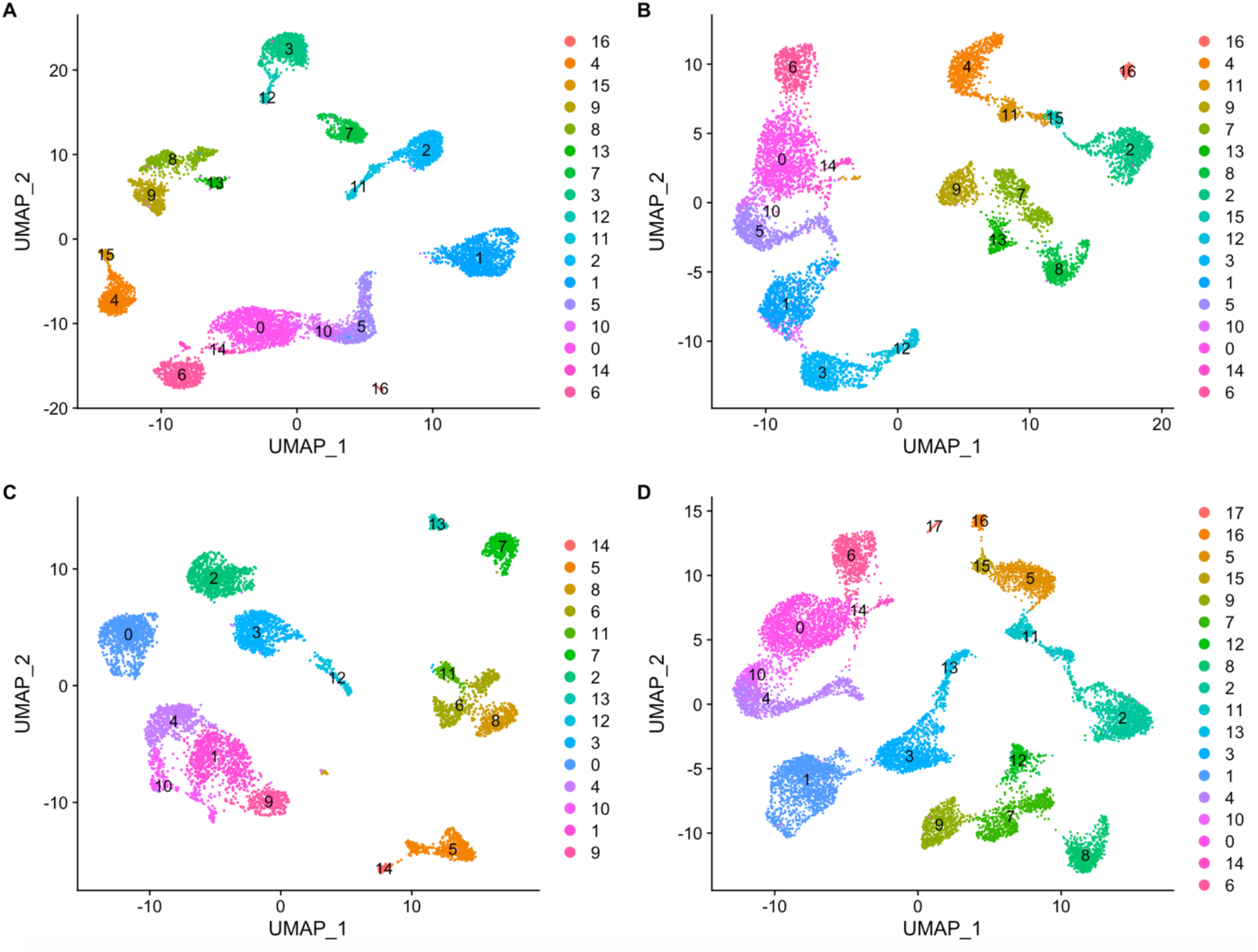
UMAP plot for different clustering comparisons. (A) Clustering of pure spercies’ testis cells samples, 3 collared flycatchers and 3 pied flycatchers. (B) Clustering of Collared flycatcher testis cell samples and hybrid samples, 3 collared flycatchers and 2 hybrid flycatchers. (C) Clustering of Pied flycatcher testis cell samples, 3 pied flycatchers and 2 hybrid flycatchers. (D) Clustering of all samples together. All the clusterings are highly consistent and vary only in the number of somatic cell cluster or in the case of panel C for the amount of clusters identified in the post meiotic stages, 5 spermatid clusters instead of the 6 spermatid clusters found in all the other clusterings.

Using the pure species reference clustering, we made projections of our 2 hybrid samples to examine equivalences with all pure species testis cell clusters (Figure 2). We find that male hybrids have all the testis cell clusters that pure species males have. However, there was a significant lower proportion of cells belonging to the spermatid clusters in hybrids (Table 1). This finding matches the histological observations of a lower overall number of spermatids produced in hybrids (Figure 3). Histology also shows an abnormal head phenotype of the developing spermatids as well as a lack of structural bundle organization of hybrid sperm cells that is observed in the samples from pure species males (Figure 3).

**Table 1.** General linear model with binomial response of spermatid cells and non-spermatid cells. We use species of our 8 individuals as explanatory variable.

| Coefficients | Estimate | Std. Error | Z value | P value | Confidence Intervals |
| --- | --- | --- | --- | --- | --- |
| Intercept | -0.37 | 0.03 | -14.5 | < **0.01** | **-0.43 - -0.32** |
| Hybrid | -0.4 | 0.05 | -8.77 | < **0.01** | **-0.49 - -0.31** |
| Pied | 0.11 | 0.04 | 2.73 | <**0.01** | **0.03 – 0.19** |

**Figure 2.**
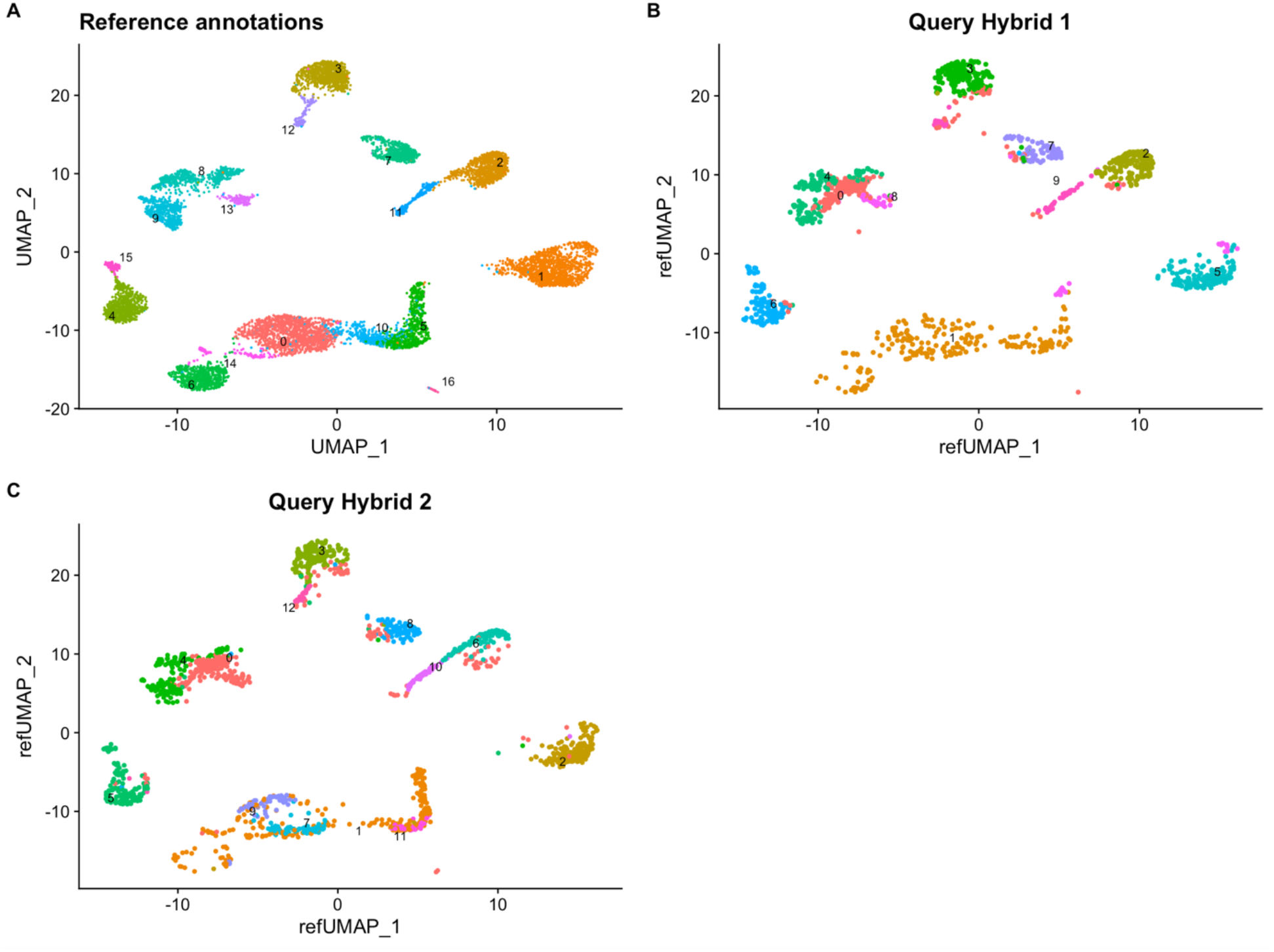
UMAP of hybrid samples mapped to the reference clustering of the pure species. Hybrid testis cells map to all stages of spermatogenesis found in the pure species, however there is a significant proportional underrepresentation of the post meiotic clusters.

**Figure 3.**
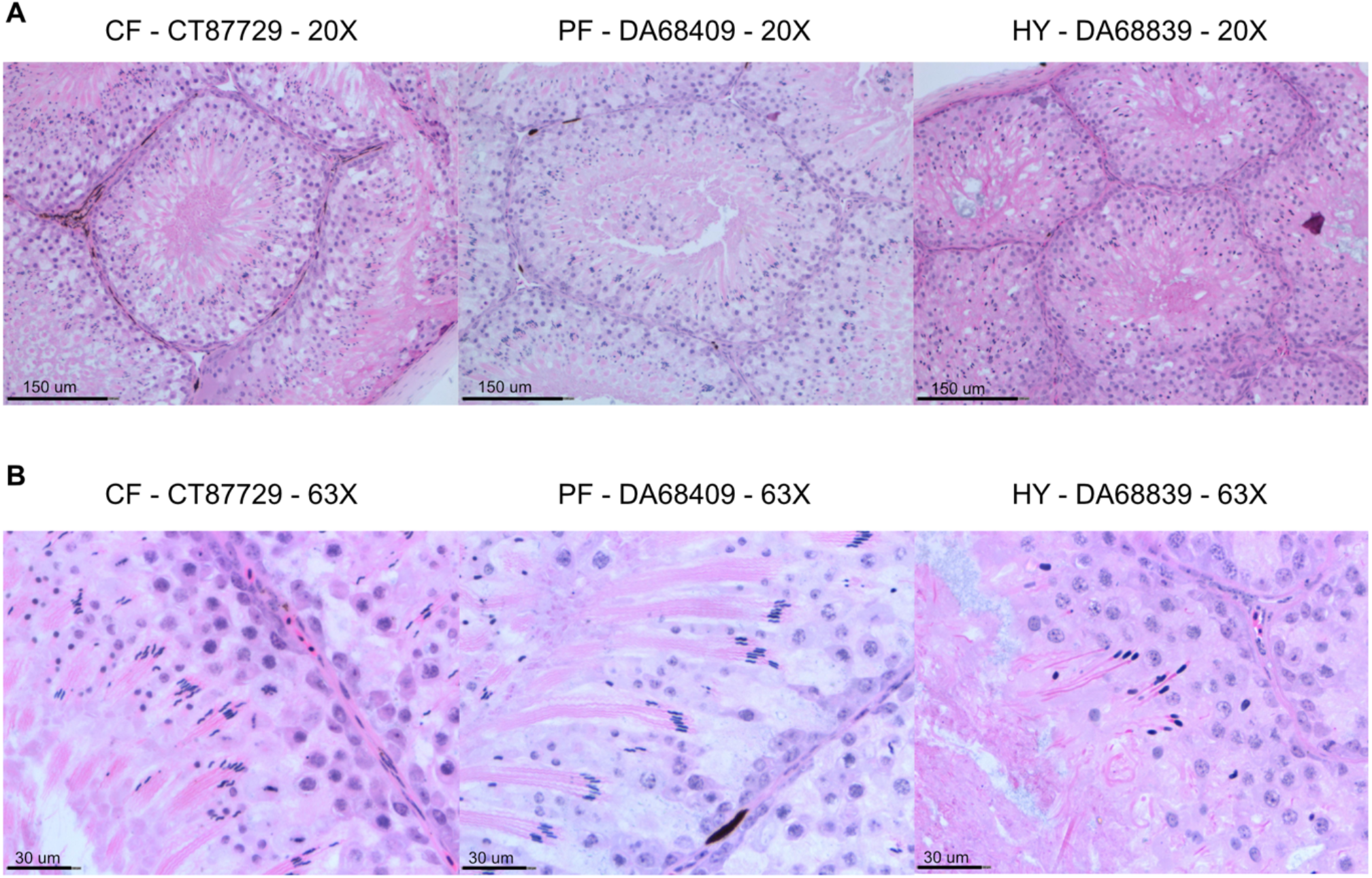
Histology sections of testis samples of Collared, Pied and Hybrids with hematoxylin and eosin staining. (A) Transversal cut of seminiferous tubuli. (B) Detail of spermatid bundles maturing towards the lumen of the tubuli. The spermatid heads have an abnormal phenotype in the hybrid and the bundle of spermatids structure is lacking. Hybrids also show less quantity of spermatids with respect to the pure species.

In order to make differential gene expression comparisons, we did three additional clustering including our hybrid samples in the following groups: collareds and hybrids, pieds and hybrids, all individuals (Figure 1B -D). We decided to perform all comparisons in case hybrids would complicate clustering due to their testis cell populations being too deviating. We found consistency across all clusters indicating that hybrids do not bias the clustering process (Figure 1). Because the IDs for equivalent cell clusters differ in the different clusterings, we assigned a common name to equivalent clusters (Figure 4).

**Figure 4.**
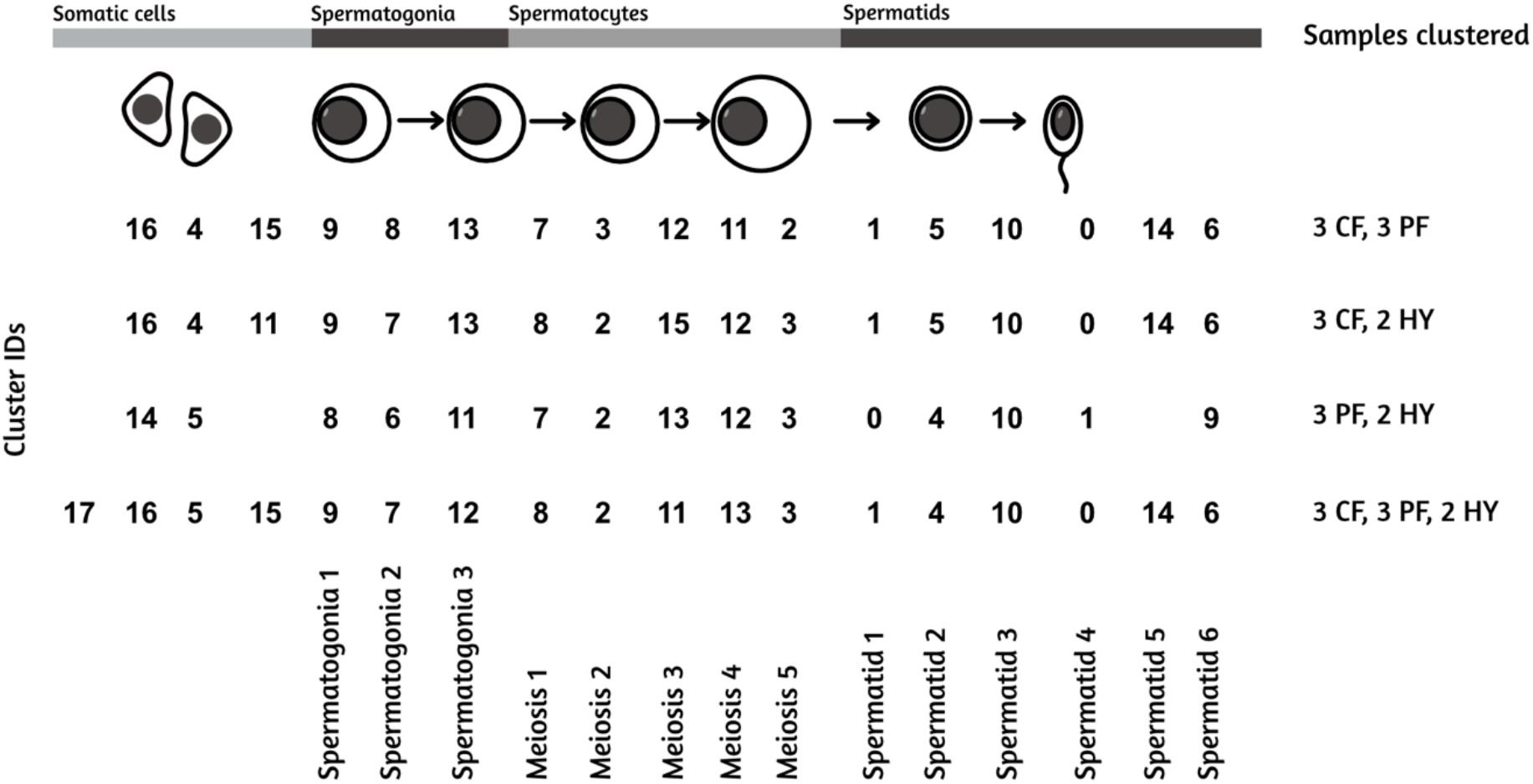
Stages of spermatogenesis and cluster IDs for the different clustering combinations.

Differential expression (DE) analysis between collared and pied, was also consistent with previous patterns found by (Segami et al 2022), most differences are found in the post-meiotic clusters, some DE genes are also found in meiosis clusters and almost no DE are found in the earliest stages of spermatogenesis (Figure 5A). The 3 comparisons of expression patterns against hybrids (i.e. collared against hybrids, pied against hybrids and pure species against hybrids) were consistent with each other. There was a slightly lower proportion of DE genes during the meiosis stages observed in the three contrasts including hybrids (Figure 5, C-D) as compared to the contrast between pied and collared flycatchers (Figure 5 A). An analysis of the hybrid inheritance patterns of gene expression shows differences between the different stages of spermatogenesis indicating that miss-expression, additive expression or a more dominant parental species expression vary along the timeline (Figure 6). For example, visual inspection suggests that “Meiosis 4” shows the strongest tendency for miss-expression (both over-dominant and under-dominant) with a narrow shape of the ellipse occupying the right top and left bottom areas of the plot. Statistical analysis is needed to assess its significance.

**Figure 5.**
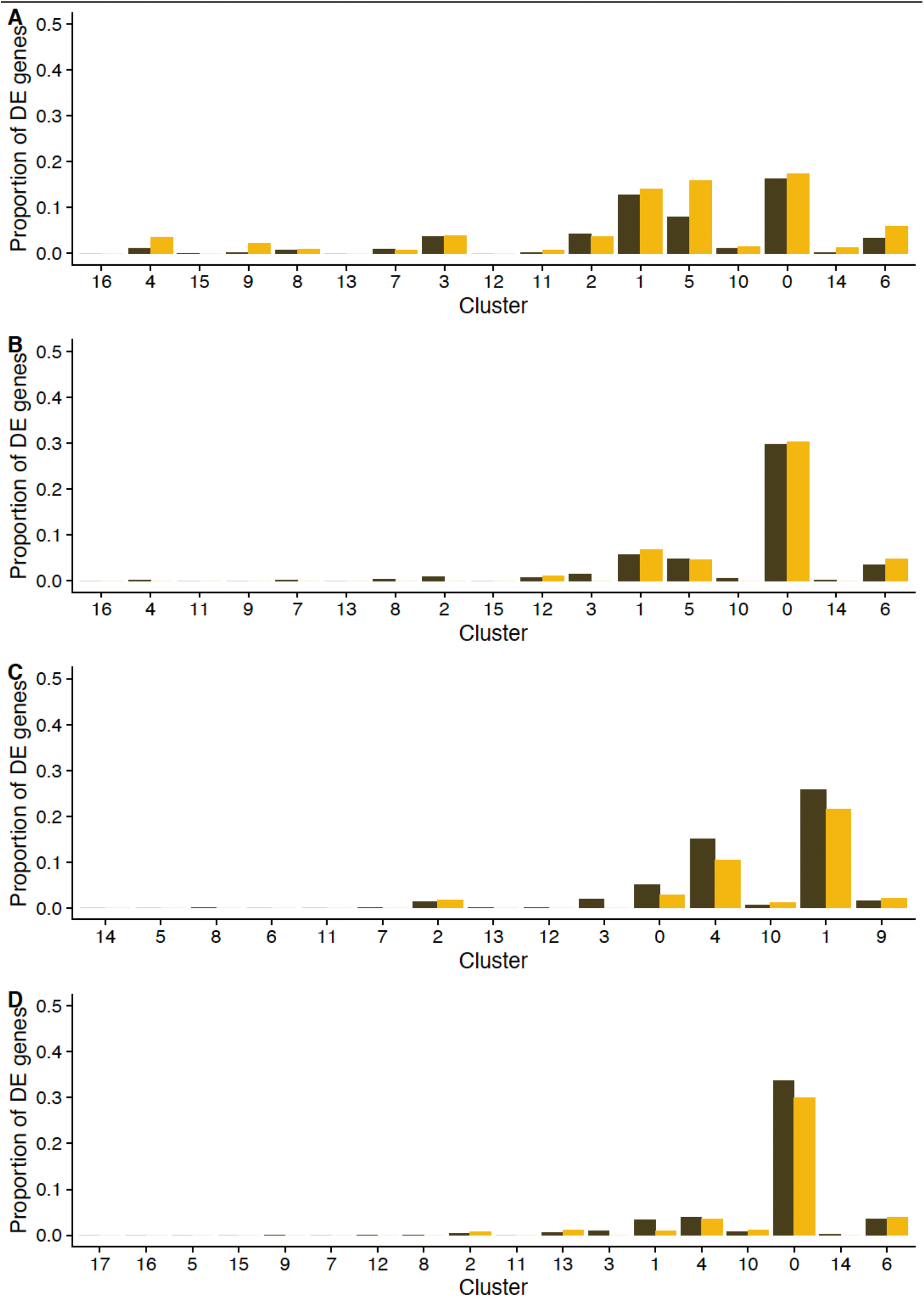
Proportion of DE genes along the timeline of cell clusters of spermatogenesis for different pairwise comparisons. (A) Collared against pied flycatcher. (B) Collared flycatcher against hybrids. (C) Pied flycatcher against hybrids. (D) Collared and pied flycatchers combined against hybrids. The general pattern of more similar gene expression at the first stage of spermatogenesis is consistent across the four comparisons. The differential expression genes start to be found in meiosis clusters and in higher proportion at the post-meiosis stages. All pairwise comparisons including hybrids show a higher proportion of DE genes than the collared vs pied comparison in one of the post meiotic clusters.

**Figure 6.**
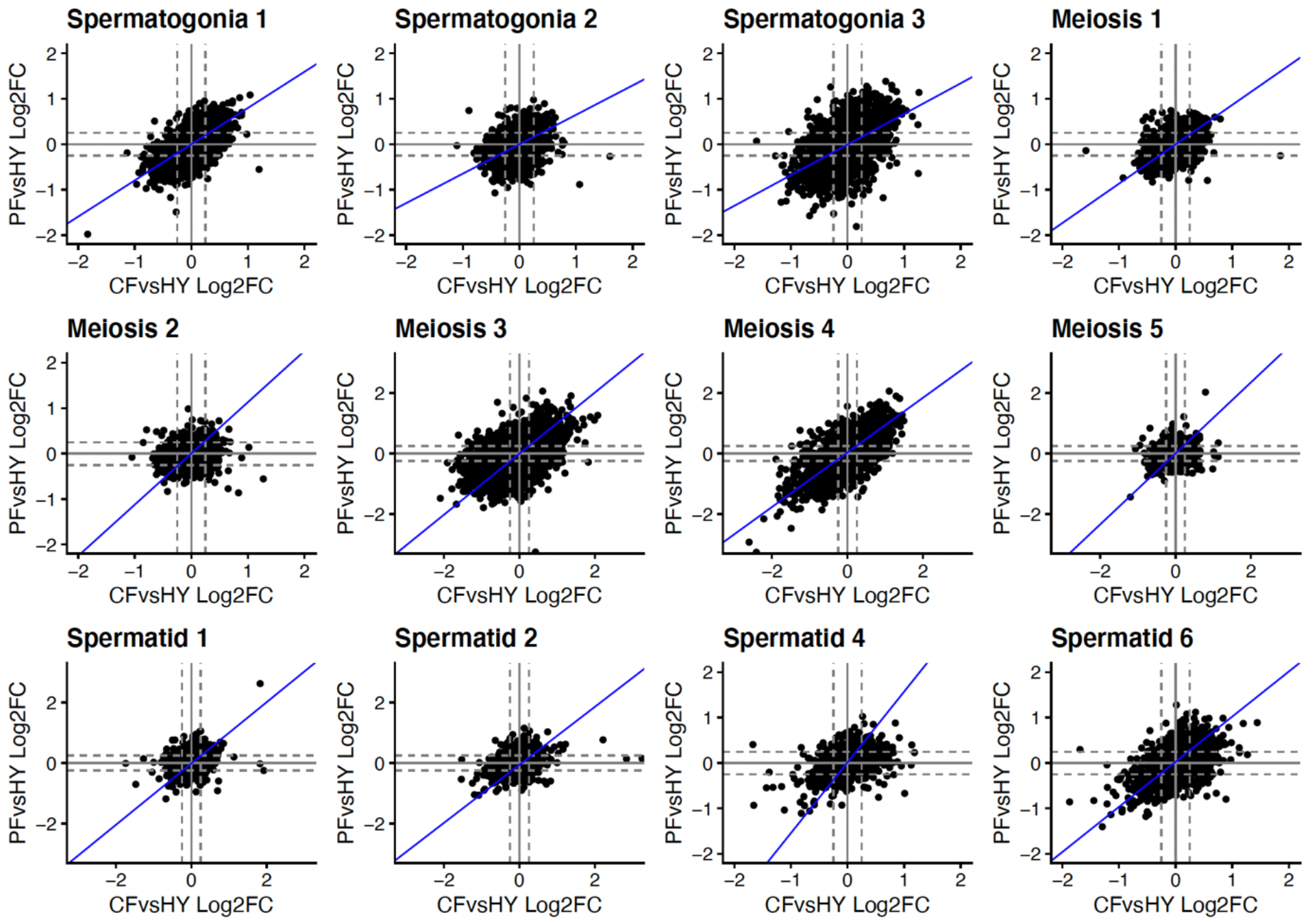
Inheritance patterns of inheritance in different stages of spermatogenesis in F1 Ficedula hybrids. Each dot represents a single gene, the grey dashed lines represent the 0.25 log2-fold change threshold, and the blue regression line is the major axis of variation in log2-fold changes. The inheritance patterns vary on the different stages of spermatogenesis, suggested by the change of the shape of the ellipsis.

We identified 303 genes with fixed non-synonymous differences between collared and pied flycatchers across the whole genome (Table S2). These genes are significantly enriched on the Z chromosome (hypergeometric test: 65,590, 21908,241, p = 2.914911e-48). Gene Ontology (GO) analysis suggest that several of these genes belong to gene networks involved in regulating several processes associated with meiosis, including DNA replication, nuclear division, chromatin remodeling and packaging (Figure 7, Table 2). Moreover, several of the genes included in the identified networks are located on the Z chromosome. We found that 71 genes with non-synonymous fixed differences between collared and pied flycatchers were expressed in the testis cell clusters (Figure 8). Interestingly, there is a clear pattern showing that they are mainly expressed in pre-meiotic and meiotic clusters, especially during Spermatogonia 1, Spermatogonia2 and Meiosis 3. By contrast, we find very few genes with fixed differences between the two species that were expressed in the spermatid clusters.

**Table 2.**
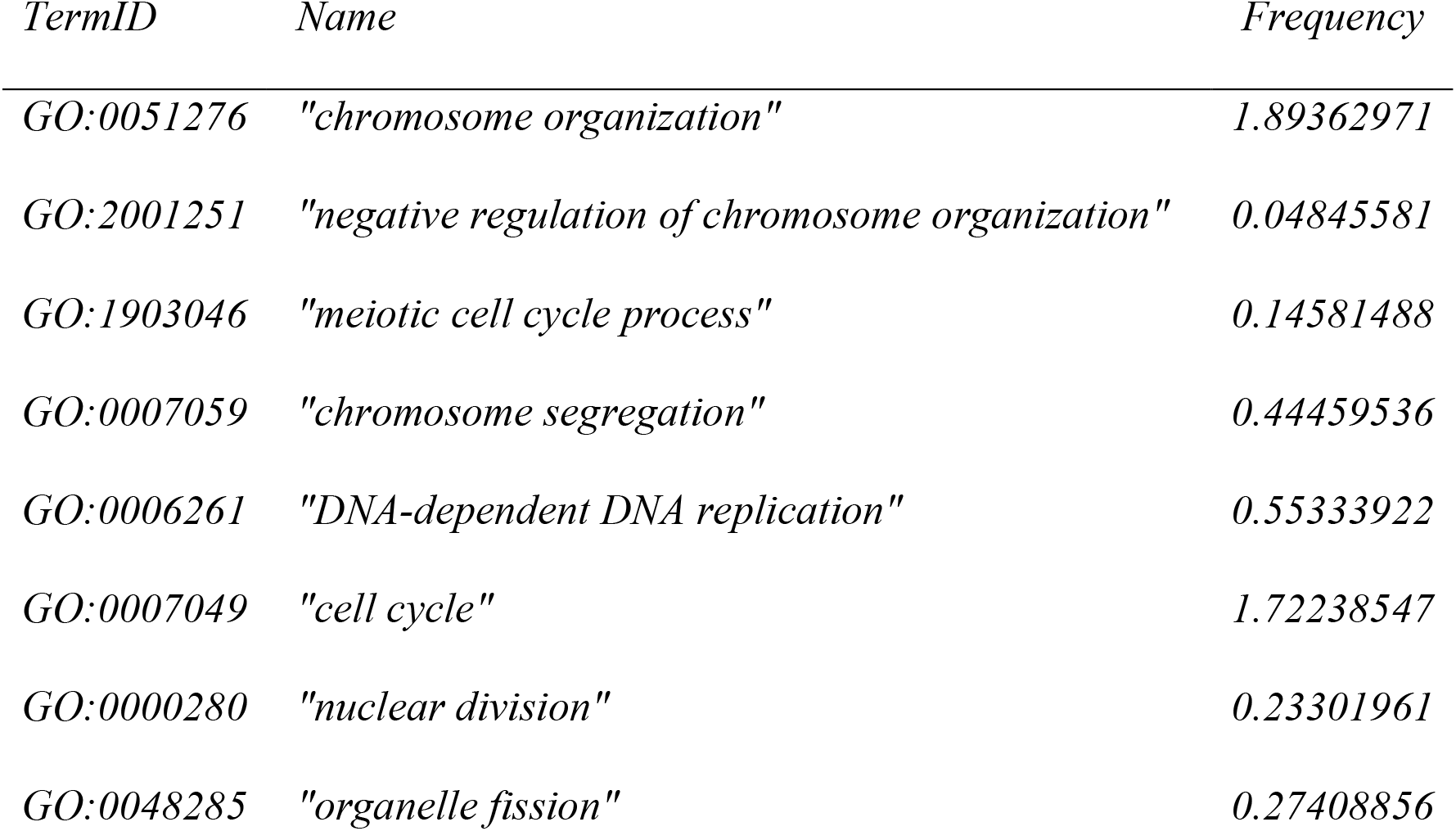
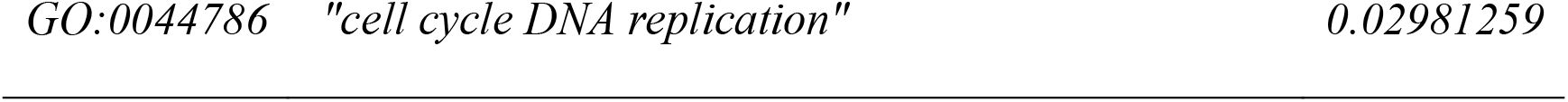
Non redundant GO terms of fixed differences analysis.

| TermID | Name | Frequency |
| --- | --- | --- |
| GO:0051276 | "chromosome organization" | 1.89362971 |
| GO:2001251 | "negative regulation of chromosome organization" | 0.04845581 |
| GO:1903046 | "meiotic cell cycle process" | 0.14581488 |
| GO:0007059 | "chromosome segregation" | 0.44459536 |
| GO:0006261 | "DNA-dependent DNA replication" | 0.55333922 |
| GO:0007049 | "cell cycle" | 1.72238547 |
| GO:0000280 | "nuclear division" | 0.23301961 |
| GO:0048285 | "organelle fission" | 0.27408856 |

**Figure 7.**
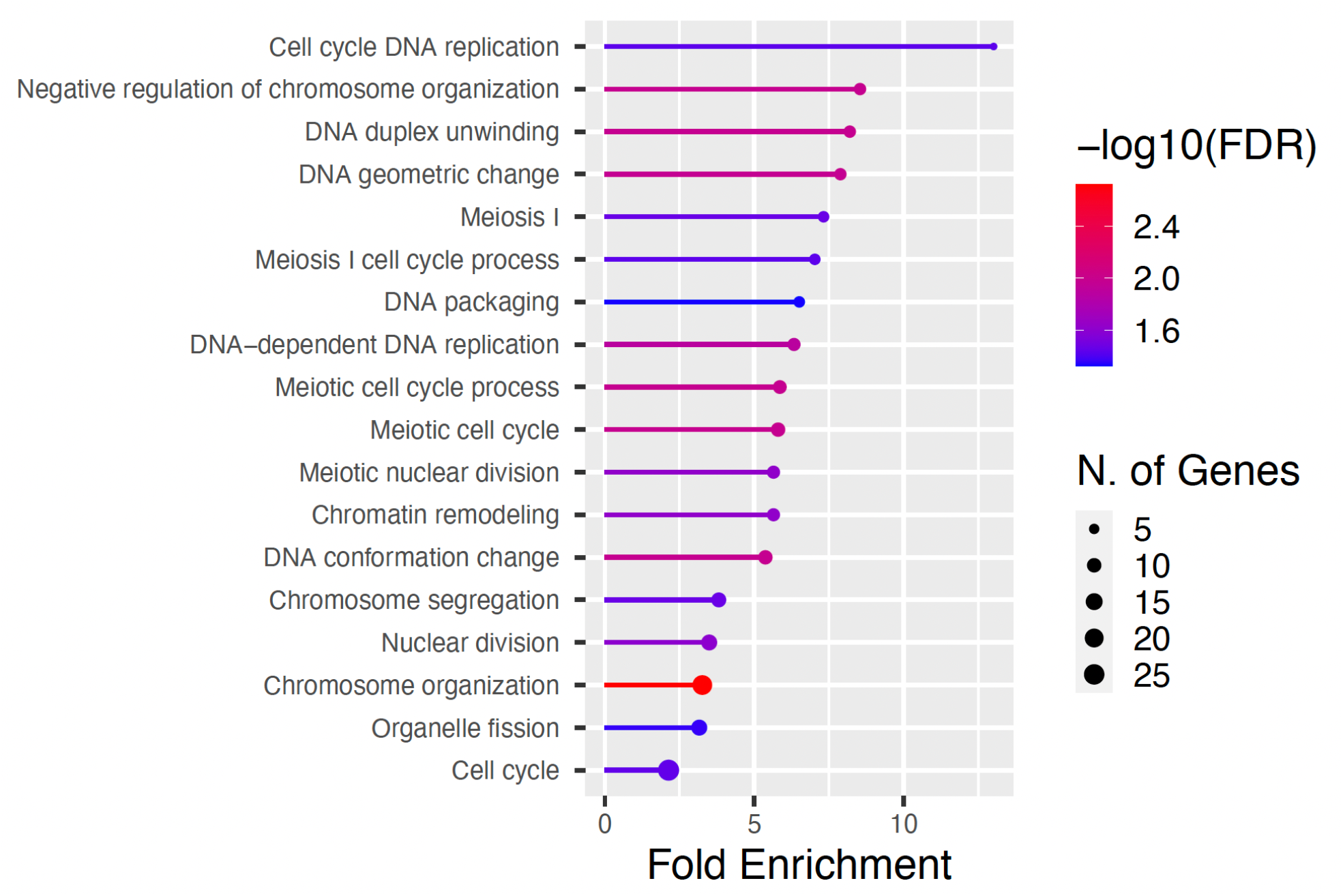
Significant GO terms for fixed non-synonymous differences between collared and pied flycatchers.

**Figure 8.**
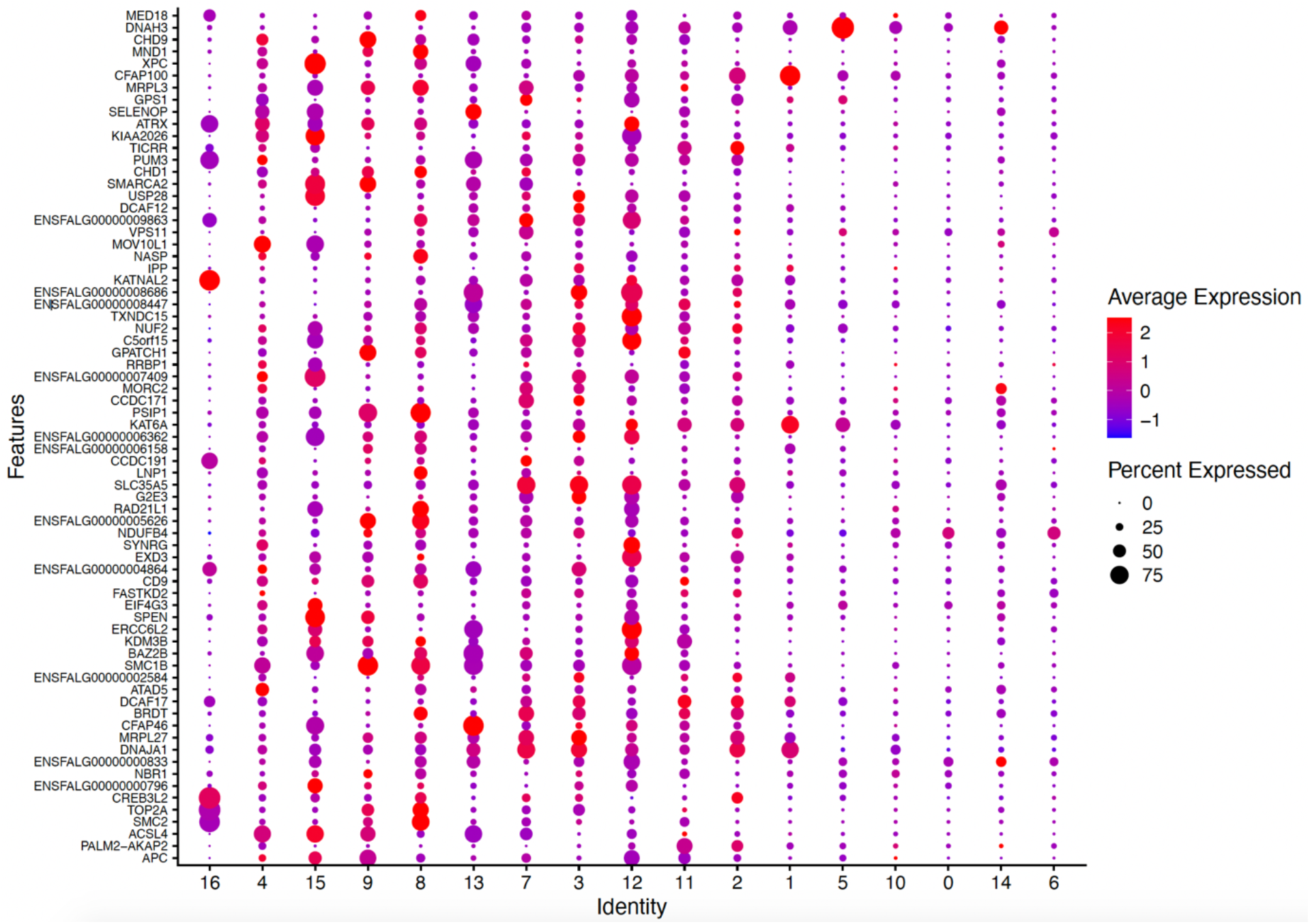
Average gene expression of genes with fixed non-synonymous differences between Collared and Pied flycatcher across the spermatogenesis stages. There is a clear pattern of absence of the fixed differences in genes expressed in spermatid clusters (1, 5, 10, 0, 14, 6). Spermatogonia and the meiosis cell clusters have the highest expression of genes with non-synonymous differences, particularly cluster 12 belonging to meiosis seems to be enriched.

A STRING analysis based on the 71 genes with non-synonymous differences that were expressed in our testis cell clusters revealed two major networks of interacting genes (Figure 9). Of these interacting genes, 3 were also DE between collared and pied flycatchers: CHD1, PSIP1 and SMC1B. CDH1 is a transcription factor associated with chromatin remodeling, PSIP1 a transcriptional co-activator involved in stem cell differentiation and SMC1B is a meiosis specific component of cohesion complex involved in chromatid movement and organization (Table 3).

**Table 3.**
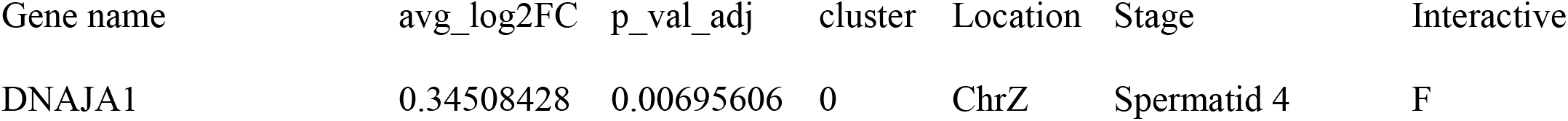
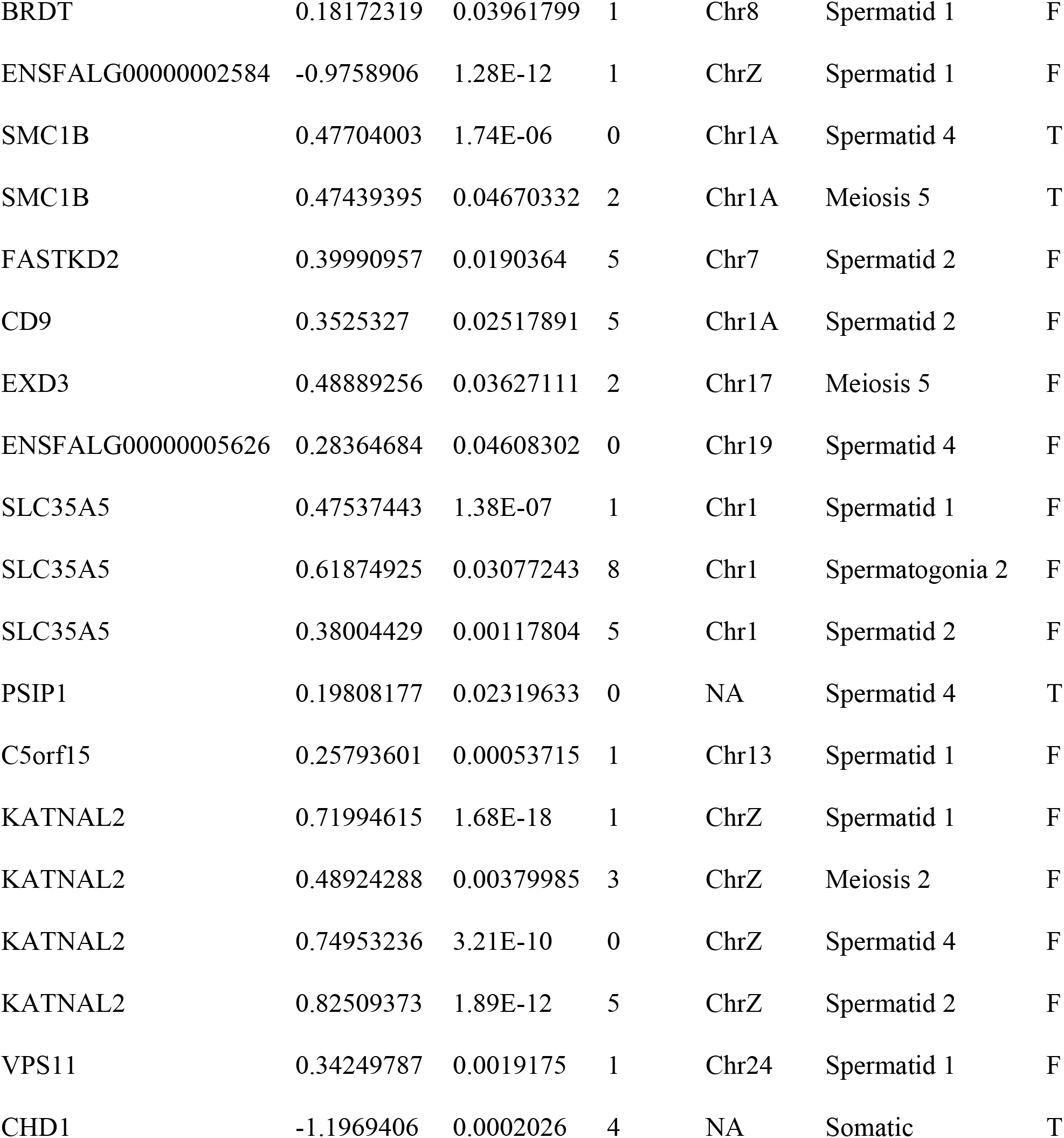
Genes with non-synonymous fixed differences between collared and pied flycatchers that are also DE.

|  |  |  |  |  |  |  |
| --- | --- | --- | --- | --- | --- | --- |
| BRDT | 0.18172319 | 0.03961799 | 1 | Chr8 | Spermatid 1 | F |
| ENSFALG00000002584 | -0.9758906 | 1.28E-12 | 1 | ChrZ | Spermatid 1 | F |
| SMC1B | 0.47704003 | 1.74E-06 | 0 | Chr1A | Spermatid 4 | T |
| SMC1B | 0.47439395 | 0.04670332 | 2 | Chr1A | Meiosis 5 | T |
| FASTKD2 | 0.39990957 | 0.0190364 | 5 | Chr7 | Spermatid 2 | F |
| CD9 | 0.3525327 | 0.02517891 | 5 | Chr1A | Spermatid 2 | F |
| EXD3 | 0.48889256 | 0.03627111 | 2 | Chr17 | Meiosis 5 | F |
| ENSFALG00000005626 | 0.28364684 | 0.04608302 | 0 | Chr19 | Spermatid 4 | F |
| SLC35A5 | 0.47537443 | 1.38E-07 | 1 | Chr1 | Spermatid 1 | F |
| SLC35A5 | 0.61874925 | 0.03077243 | 8 | Chr1 | Spermatogonia 2 | F |
| SLC35A5 | 0.38004429 | 0.00117804 | 5 | Chr1 | Spermatid 2 | F |
| PSIP1 | 0.19808177 | 0.02319633 | 0 | NA | Spermatid 4 | T |
| C5orf15 | 0.25793601 | 0.00053715 | 1 | Chr13 | Spermatid 1 | F |
| KATNAL2 | 0.71994615 | 1.68E-18 | 1 | ChrZ | Spermatid 1 | F |
| KATNAL2 | 0.48924288 | 0.00379985 | 3 | ChrZ | Meiosis 2 | F |
| KATNAL2 | 0.74953236 | 3.21E-10 | 0 | ChrZ | Spermatid 4 | F |
| KATNAL2 | 0.82509373 | 1.89E-12 | 5 | ChrZ | Spermatid 2 | F |
| VPS11 | 0.34249787 | 0.0019175 | 1 | Chr24 | Spermatid 1 | F |
| CHD1 | -1.1969406 | 0.0002026 | 4 | NA | Somatic | T |

**Figure 9.**
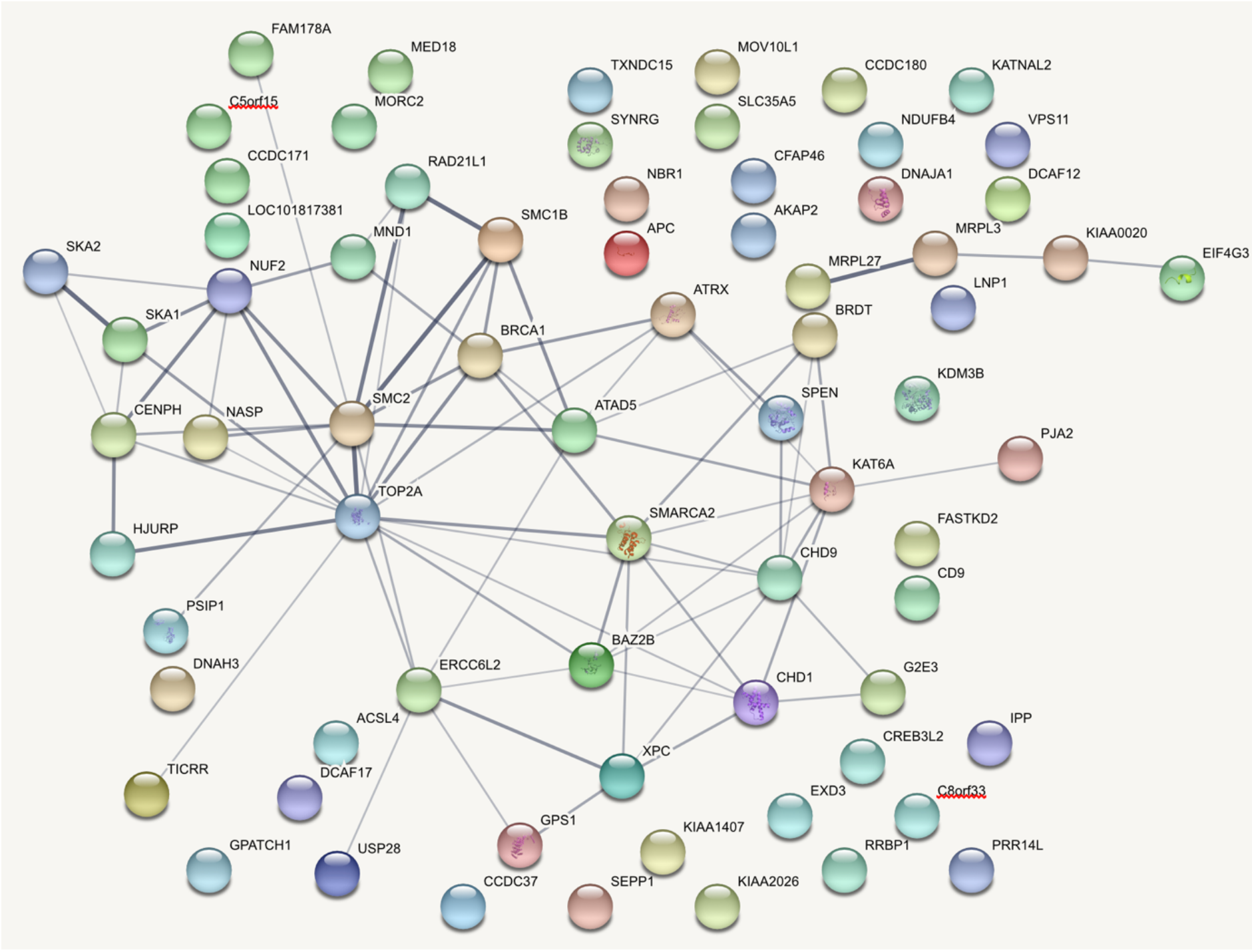
STRING Interacting gene networks for genes with non-synonymous fixed differences that are expressed in our spermatogenesis cell populations.

We also found that some of the non-interacting genes were DE (Table 3). Of these, EXD3, SLC35A5 and KATNAL2 are DE in spermatogonia and meiosis cell clusters. EXD3 is an exonuclease, SLC35A5 is transmembrane sugar transporter and KATNAL2 is a katanin microtubule-severing protein.

## Discussion

Identifying genes involved in genetic incompatibilities causing hybrid sterility or inviability is a long-standing challenge in speciation research. Here we present several lines of evidence implying that hybrid male sterility in *Ficedula* flycatchers is associated with a failure of meiosis and we propose candidate genes involved in genetic incompatibilities causing this failure. We conclude that dysfunctional meiosis most likely leads to aberrant chromosome segregation and/or faulty chromatin packing, similarly to what has been found in mouse (Mihola et al., 2009) and drosophila (Kanippayoor et al., 2020). This conclusion is based on combined evidence from whole genome DNA re-sequencing data, single cell transcriptomics of testis samples and testis histology sections. STRING and GO analysis show that genes with non-synonymous differences between the two species of flycatchers are part of gene networks involved in key processes of meiosis. A lower proportion of spermatids produced by hybrid males as revealed by histology and single cell data implies that a proportion of the aberrant spermatids undergo apoptosis.

Based on the histological observations of a lower overall number of spermatids, all with an abnormal head phenotype and lacking the structural bundle organization in the testis of hybrid flycatchers (Figure 3), we were surprised to find that all main cell clusters were present in their testis samples. However, many genes with non-synonymous fixed differences between collared and pied flycatchers were expressed during, the early and most fundamental developmental stage of spermatogenesis (i.e. during Spermatogonia 1, Spermatogonia2 and Meiosis 3). Moreover, we revealed two major networks of interacting genes (Figure 10) where 3 genes stand-out by also being DE between collared and pied flycatchers: CHD1, PSIP1 and SMC1B. In addition, we detected three additional genes with fixed-differences that were DE in spermatogonia and meiosis cell clusters: EXD3, SLC35A5 and KATNAL2. Based on the known central functions of all these genes (see Table 3) failure during meiosis is a highly likely explanation for dysfunction of spermatogenesis in hybrid *Ficedula* flycatchers.

Two of the BDMI candidate genes for hybrid sterility in flycatchers have been described before as having a crucial role during spermatogenesis and/or for maintaining fertility. SCMB1 is required for sister chromatid cohesion and DNA recombination in mouse and deficient SCMB1 mice of both sexes show sterility associated to pachytene in males and meiosis II failure in females (Revenkova et al., 2004). This roughly correspond with the stage where we observe meiosis failure in flycatchers, and although this study focuses only on male gametogenesis, we know that female F1 hybrids are also sterile as they produce eggs that do not hatch(Svedin et al., 2008). It is also shown that a spontaneous mutation in the SCMB1 mice gene caused sterility in both sexes (Takabayashi et al., 2009). In addition, SCMB1 also causes sterility in both sexes of zebrafish impairing correct synapsis (Islam et al., 2021).

Thus, although SCMB1 has a slightly different role among vertebrates, this gene is essential for correct meiosis. One of the most interesting things is that in all these cases there was an essential role of SCMB1 for fertility but not for sperm development, i.e. spermatogenesis was not arrested but sperm cells were not fertile, this aligns perfectly with our observations in flycatchers. KATNAL2 has several essential functions related to microtubule dynamics, cytokinesis and ciliogenesis. Knockouts of this gene results in aberrant phenotypes of cells showing multinuclearity and abnormal sperm heads in mice (Ververis et al., 2016; Houston et al., 2020). This gene also has a critical role during spermiogenesis as it is involved in sperm tail growth, sperm head shaping, acrosome attachment and sperm release to the lumen of seminiferous tubuli (Dunleavy et al., 2017). The defects described for mice caused by mutations or knockout of KATNAL2 fit well with our observation of abnormal head shape in the histology samples of the testis of hybrid flycatchers.

Our results may appear contractionary in the light of the first developmental stages of spermatogenesis (i.e. mitosis and meiosis) being known to be strongly evolutionary constrained both in mammals (Larson et al., 2018; Kopania et al., 2022) and in birds (Segami et al 2022). Most of the known examples of genes associated with hybrid sterility in animals are, in fact, expressed during either the pre-or post-meiotic stages of gametogenesis (Coyne & Orr, 2004). However, most of these examples come from studies based on drosophila species that usually have diverged for several millions of years. The only previously known example of a specific gene causing hybrid sterility in vertebrates is based on studies of a natural hybrid zone between two closely related species of mouse. In that case, *prdm9* was shown to cause hybrid sterility through disrupted meiosis (Mihola et al., 2009). One possible explanation to the findings in flycatchers and mice is that disruption of meiosis in hybrids quickly leads to reproductive isolation, which in turn leads to accumulation of genetic differences in faster evolving regions of the genome that may rather be a consequence than a cause of speciation.

In agreement with both theoretical expectations and empirical findings based on study species with XY sex-determination systems, our findings imply an important role of Z-linked BDMIs in speciation. The genes expressed during spermatogenesis that contain non-synonymous differences between the two flycatcher species are enriched on the Z chromosome and three of our BDMI candidate genes are Z-linked. Two additional detected candidate genes, CHD1 and PSIP1 are located on a scaffold that remains unassigned in the flycatchers but are known to be located on the Z chromosome in the zebra finch. Since birds have a relatively stable karyotypes which also tend to keep gene order across species (Ellegren, 2010), we consider it highly likely that these genes are Z-linked also in flycatchers. This remains to be confirmed once a better assembly of the flycatcher genome is available.

The latest evidence for fast Z, not only in flycatchers (Ellegren et al., 2012; Mugal et al., 2020) but also in birds in general, points to drift rather than selection (Mank et al., 2010) as a main driving evolutionary force. However, since drift is a very slow evolutionary process, we consider drift unlikely to explain species divergence in a highly evolutionary conserved function such as meiosis. Instead, selection may have acted on standing genetic variation at the very early stages of the split between the two flycatchers resulting in comparatively fast lineage sorting of ancestral allelic variation in these particular genes.

Our findings correspond well with what has been previously described in model species with XY-sex determination systems. The two proteins encoded by SCMB1 and KATNAL2 are with a high probability central for causing genetic incompatibilities in F1 *Ficedula* hybrids. Since a handful of incompatibilities, rather than a simply two allele single incompatibility contribute to sterility in hybrid flycatchers, several incompatibilities may have arisen in a snowballing fashion. Although milder incompatibilities between fast evolving alleles are predicted to arise first during genetic divergence our results points in the opposite direction with the most serious incompatibilities affecting central functions having arisen first. This could be due to meiosis being a particularly sensitive process given the complicated networks of genes. One change could then cause selection favoring compensatory changes in epistatically interacting genes thereby unleashing snowballing of incompatibilities.

This study is to our knowledge the first one to propose candidate BDMI genes causing hybrid sterility in birds and together with (Rosser et al., 2022) the second study to propose candidate genes causing hybrid sterility in ZW systems. We revealed evidence for disrupted meiosis during spermatogenesis and an overrepresentation of Z-linkage of the targeted candidate incompatibility genes. Our results challenge the assumption that speciation processes are driven by fast evolving genes by showing that changes in genes with highly conserved functions may, when slightly modified, be likely to quickly cause speciation by ensuring reproductive isolation at secondary contact.

## Methods

### Samples and sequencing

Three collared flycatchers, three pied flycatcher and two F1 hybrids with a pied mother were caught at the start of the reproductive season in the monitored populations of Öland (57°100N, 16°580E) (Anna Qvarnström et al., 2010), Sweden in May 2020. The birds were kept in outdoor aviaries until all birds were caught before being sacrificed by cervical dislocation and immediately dissected. The testes were cleaned from other tissues and cut in halves on a cold petri dish with PBD BSA buffer. A half of testis was mechanically disassociated in 3ml of PBD BSA buffer using a gentleMACS Dissociator (Miltenyi Biotech, Bergisch Gladbach, Germany) as described in (Segami et al 2022). We verified that cell viability was over 80% by examining the cell suspensions under the microscope with propidium iodide and Hoechst staining. Then the cell suspension concentration was determined with a Neubauer chamber and diluted to achieve 1×10^6 cell/ml. The final cell suspensions were delivered to the sequencing platform for library preparations with 10X genomics Chromium Single Cell 3’ v3 kit for scRNAseq. All the libraries were sequenced in two lanes of NovaSeq S1 flow cells.

### Data pre-processing

We used the 10x Genomics Cell Ranger v. 6.0.0 (Zheng et al., 2017) pipeline, we created a custom reference for cell counting based on the public genome for *Ficedula albicollis* (v. 1.4) and the public annotation from *Ensembl* (v. 1.4). After obtaining count matrixes for every gene on each cell for our 8 samples, we generated .loom files for each individual using the command run10X from the python package Velocyto v. 0.17.17 (La Manno et al., 2018). Finally, we exported all the loom files to Seurat v. 4.1.0 (Stuart et al., 2019) for downstream analysis.

### Filtering, normalization, and clustering

We converted all the loom files to Seurat objects and merged the objects in the following combinations: only pure species (6 individuals), Collareds and hybrids (5 individuals), Pied and hybrids (5 individuals), all individuals (8 individuals). We excluded all cells with less than 200 features (expressed genes), more than 2500 and less than 5% of mitochondrial genes. For normalization, variance stabilization and integration of the different samples in a single object we used the SCTransform v2 (Choudhary & Satija, 2022) command together with anchor integration as suggested on the Seurat documentation. Then we performed standard clustering analysis for all the combinations with the following parameters, we run UMAP (Uniform Manifold Approximation and Projection) with 20 dimentions and min.dist = 0.6. Finally, we found clusters using a resolution of 0.5. For all the 4 clusterings we found an heterogeneous cluster with no good markers that we excluded and reclustered the remaining cells again. After obtaining all our final clusterings we established the identity of all clusters using the flycatcher gene markers characterized by (Segami et al 2022). We then established the equivalences of the clusters across our different clusterings (Figure 4).

### Histology

Whole testes were preserved in formalin for 3 collareds, 3 pieds and 2 hybrids. Histology sections were done for each sample fixed to slides and then stained with hematoxylin and eosin. Pictures were taken using an optical microscope with magnification 65X and 20X.

### Hybrid cell clusters analysis

In order to test if our hybrid samples contained all the cell clusters present in the spermatogenesis of pure species, we performed a unimodal UMAP projection of each hybrid sample on the reference clustering of the pure species. We visually observed in the clustering analysis that the proportion of spermatids was lower in both hybrids and therefore decided to test whether the proportion of spermatids was significantly different than the proportion of spermatids found in pure species males. For this purpose, we clustered all our samples individually and identified the proportion of total cells that corresponded to spermatids. Then we performed a binomial general linear model with number of spermatid cells and non-spermatid cells as response variable and species as explanatory variable.

### Differential expression analysis

We performed DE analysis for all our clusterings comparing collared vs. pied, collared vs. hybrid, pied vs. hybrid and pure species vs. hybrid. The Seurat function FindMarkers was used to perform a Wilcoxon rank sum test to calculate average fold change and identify DE genes. We used the normalized counts from RNA slot of our Seurat objects and did not use a minimum threshold for fold change. To establish if a gene was significantly DE, we used the adjusted p-value with a 0.05 threshold. In order to explore the hybrid gene expression inheritance patterns for each cluster, we plotted the obtained fold changes for the comparisons of collared vs. hybrid and pied vs. hybrid for each gene. We then calculated the major variation regression line using the r package lmodel2.

### Identification of non-synonymous fixed differences

Fixed differences between collared flycatcher and pied flycatcher were identified based on polymorphism data for 19 collared and 19 pied flycatchers from the island of Öland retrieved from (Burri et al., 2015). We excluded polymorphic sites with more than two alleles at a specific site and further restricted the data to sites that were covered by at least 1 read across all 38 individuals. Sites that were classified as fixed differences if all collared and pied individuals showed a different allele. Fixed differences were annotated using SnpEff version 4.3T (Cingolani et al., 2012) based on Ensembl gene annotation v 73 (Uebbing et al., 2016), which resulted in a set of 303 non-synonymous fixed differences across 21908 genes.

### Gene Ontology and STRING Analysis

We did Gene Ontology enrichment analysis for biological processes on the set of genes with non-synonymous fixed differences and on the subset of these genes that were expressed in our testis cell clusters using the web tool ShinyGO v.075 (Ge et al., 2020). To prescind of the redundant and/or nested GO terms we used the web tool REVIGO (Supek et al., 2011). To identify the interacting gene networks we performed STRING (Szklarczyk et al., 2019) analysis using their web tool v.11.5 on the same sets of genes.

## Author Contributions

AQ and JCS conceived the study. AQ, JCS and CC collected the samples. JCS sacrificed the birds. CC performed the dissections. JCS, CC and CB carried out the cell suspension protocol. MSz performed the histology. JCS and MS performed the single cell clustering and bioinformatics analysis. CFM performed bioinformatics analysis and identified the fixed differences. JCS, MS and CFM performed and discussed the GO analysis and all the statistical analysis. JCS, MS, CFM and AQ discussed and interpreted all the results. JCS and AQ wrote the manuscript.

## Acknowledgements

We are grateful for help during the field work or data analysis to Tom van der Valk, William Jones, David Wheatcroft, Javier Florenza García, Alexander Suh, Francisco Ruiz Ruano, Axel Jensen and all Qvarnström lab members and field assistants. We also thank the Microbial Single Cell Genomics Facility at SciLifeLab for letting us use their lab facilities. Sequencing was performed by the SNP&SEQ Technology Platform in Uppsala. The facility is part of the National Genomics Infrastructure (NGI) Sweden and Science for life Laboratory. The SNP&SEQ Platform is also supported by the Swedish Research Council and the Knut and Alice Wallenberg Foundation. Computations were performed on resources provided by the Swedish National Infrastructure for Computing (SNIC) at Uppsala Multidisciplinary Center for Advanced Computational Science (UPPMAX). Funding: This work was supported by the Swedish Research Council, grant number 2016–05138 and 2012– 03722 to AQ. CFM is funded by grants to Hans Ellegren from the Swedish Research Council (2013/08271) and Knut and Alice Wallenberg Foundation. MS is funded by IUF (Institut Universitaire de France) and ANR 21 T-ERC PLEIOTROPY and received a stipend from Wenner-Gren for a sabbatical at Uppsala University.

## Ethical permits

Permit for keeping flycatchers in aviaries and sacrificing maximum 17 flycatchers per year. Swedish environmental protection agency Natur vårds verket (NV-02622-20) valid from 2020-05-01 till 2021-06-30.

