## Supplementary materials for "The genomic basis of hybrid male sterility in *Ficedula* flycatchers"

The supplementary material includes: Figure S1 and Table S1-2.

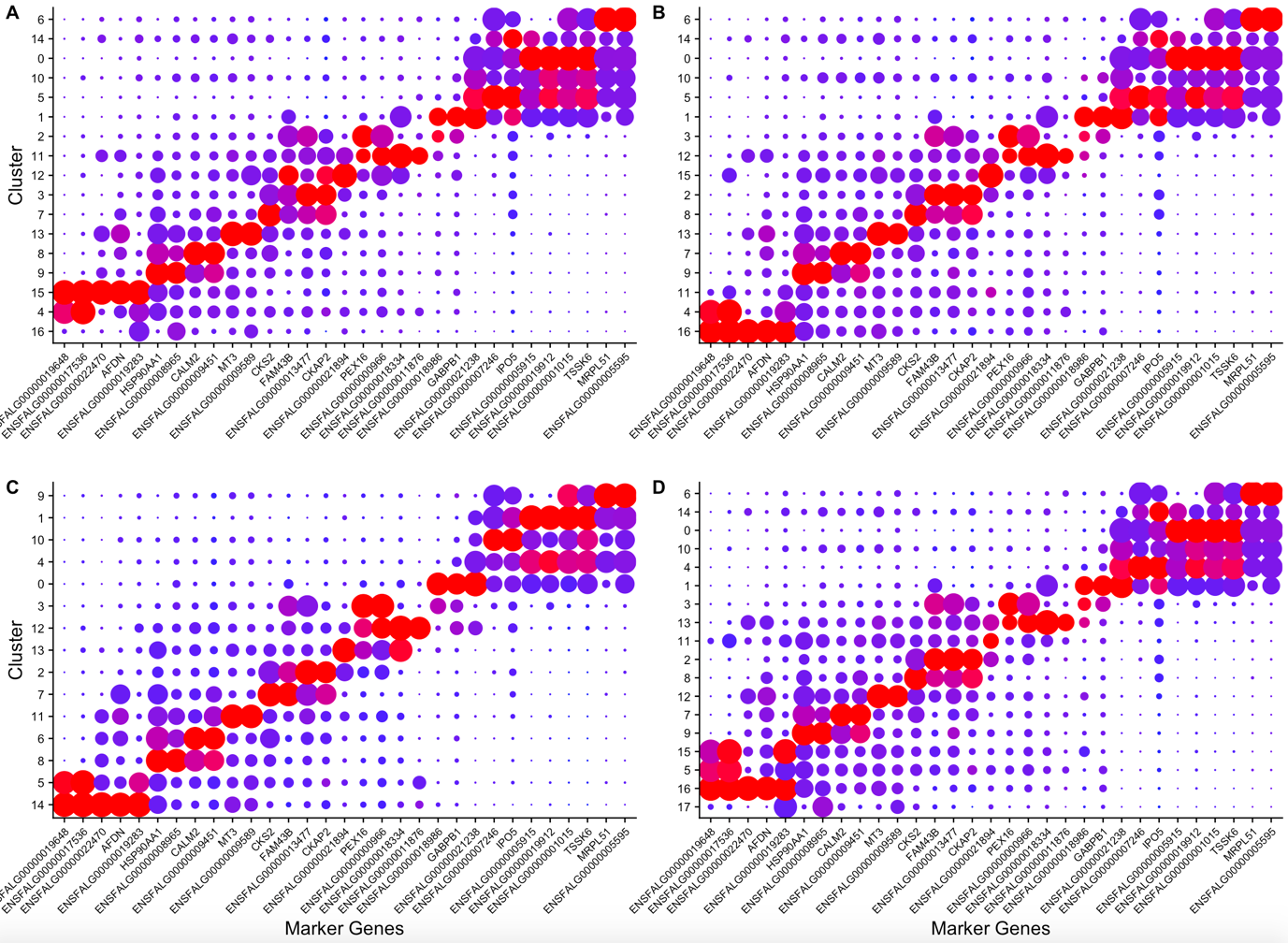

Figure S1. Average expression of flycatcher spermatogenesis markers in the different clusters identified. (A) Pure species. (B) Collared flycatcher and hybrid individuals. (C) Pied flycatcher and hybrid individuals. (D) All individuals.

Table S1. Number of total recovered cells for each sample after pre-processing and total number of cells belonging to the spermatid stage.

| Spermatid cell # | Total cells # | ID | Species | Spermatid prop | Non spermatid cell # |
| --- | --- | --- | --- | --- | --- |
| 486 | 1033 | B1 | PF | 0.47047435 | 547 |
| 858 | 1816 | B2 | CF | 0.47246696 | 958 |
| 539 | 1395 | B3 | PF | 0.38637993 | 856 |
| 931 | 2080 | B4 | PF | 0.44759615 | 1149 |
| 1054 | 2845 | B5 | CF | 0.37047452 | 1791 |
| 425 | 1581 | B6 | HY | 0.2688172 | 1156 |
| 614 | 1709 | B7 | HY | 0.35927443 | 1095 |
| 624 | 1561 | B8 | CF | 0.39974375 | 937 |

Table S2. Genes with non-synonymous fixed differences.

| Ensembl Gene ID | Symbol | Gene Type | Chromosome |
| --- | --- | --- | --- |
| ENSFALG00000005183 |  | protein_coding | 1 |
| ENSFALG00000008260 |  | protein_coding | 1 |
| ENSFALG00000008748 | ZMYM2 | protein_coding | 1 |
| ENSFALG00000005312 | NDUFB4 | protein_coding | 1 |
| ENSFALG00000005370 |  | protein_coding | 1 |
| ENSFALG00000005402 | PLA1A | protein_coding | 1 |
| ENSFALG00000005793 | F5 | protein_coding | 1 |
| ENSFALG00000005838 | SLC35A5 | protein_coding | 1 |
| ENSFALG00000005849 |  | protein_coding | 1 |
| ENSFALG00000005862 |  | protein_coding | 1 |
| ENSFALG00000005909 |  | protein_coding | 1 |
| ENSFALG00000005992 | CCDC191 | protein_coding | 1 |
| ENSFALG00000005929 |  | protein_coding | 1 |
| ENSFALG00000006362 |  | protein_coding | 1 |
| ENSFALG00000001488 |  | protein_coding | 1 |
| ENSFALG00000007434 |  | protein_coding | 2 |
| ENSFALG00000009096 | METTL6 | protein_coding | 2 |
| ENSFALG00000014345 | XCR1 | protein_coding | 2 |
| ENSFALG00000012372 |  | protein_coding | 2 |
| ENSFALG00000002146 | TERT | protein_coding | 2 |
| ENSFALG00000005911 | MEP1B | protein_coding | 2 |
| ENSFALG00000011214 | PNPT1 | protein_coding | 3 |
| ENSFALG00000013661 | BFSP1 | protein_coding | 3 |
| ENSFALG00000013666 |  | protein_coding | 3 |
| ENSFALG00000007560 | RRBP1 | protein_coding | 3 |
| ENSFALG00000007607 | DZANK1 | protein_coding | 3 |
| ENSFALG00000011587 | CNST | protein_coding | 3 |
| ENSFALG00000012083 | LRRC73 | protein_coding | 3 |
| ENSFALG00000012173 | PRKN | protein_coding | 3 |
| ENSFALG00000003388 | GALNT2 | protein_coding | 3 |
| ENSFALG00000012671 | PLAGL1 | protein_coding | 3 |
| ENSFALG00000012397 | FBXO5 | protein_coding | 3 |
| ENSFALG00000011511 | ECHDC1 | protein_coding | 3 |
| ENSFALG00000001698 | APOB | protein_coding | 3 |
| ENSFALG00000001655 |  | protein_coding | 3 |
| ENSFALG00000001547 |  | protein_coding | 3 |
| ENSFALG00000000934 |  | protein_coding | 4 |
| ENSFALG00000006605 |  | protein_coding | 4 |
| ENSFALG00000014741 |  | protein_coding | 4 |
| ENSFALG00000010624 | PARVA | protein_coding | 5 |
| ENSFALG00000010607 | MICAL2 | protein_coding | 5 |
| ENSFALG00000001067 | LUZP2 | protein_coding | 5 |
| ENSFALG00000012137 | LRRC56 | protein_coding | 5 |
| ENSFALG00000005783 | G2E3 | protein_coding | 5 |
| ENSFALG00000014925 |  | protein_coding | 5 |
| ENSFALG00000006948 | SAMD8 | protein_coding | 6 |
| ENSFALG00000007971 |  | protein_coding | 6 |
| ENSFALG00000008686 |  | protein_coding | 6 |
| ENSFALG00000008772 |  | protein_coding | 6 |
| ENSFALG00000001323 |  | protein_coding | 6 |
| ENSFALG00000001338 |  | protein_coding | 6 |
| ENSFALG00000004122 | TRAK2 | protein_coding | 7 |
| ENSFALG00000003964 | FASTKD2 | protein_coding | 7 |
| ENSFALG00000001985 | DCAF17 | protein_coding | 7 |
| ENSFALG00000004359 |  | protein_coding | 7 |
| ENSFALG00000004041 | SLC11A1 | protein_coding | 7 |
| ENSFALG00000013550 |  | protein_coding | 7 |
| ENSFALG00000003242 |  | protein_coding | 7 |
| ENSFALG00000008447 |  | protein_coding | 8 |
| ENSFALG00000007941 | NUF2 | protein_coding | 8 |
| ENSFALG00000009528 | MAST2 | protein_coding | 8 |
| ENSFALG00000009544 | IPP | protein_coding | 8 |
| ENSFALG00000009547 | TMEM69 | protein_coding | 8 |
| ENSFALG00000009600 | AKR1A1 | protein_coding | 8 |
| ENSFALG00000009566 | NASP | protein_coding | 8 |
| ENSFALG00000005047 |  | protein_coding | 8 |
| ENSFALG00000001764 | BRDT | protein_coding | 8 |
| ENSFALG00000007789 |  | protein_coding | 9 |
| ENSFALG00000005352 | HPS3 | protein_coding | 9 |
| ENSFALG00000010716 | SCAPER | protein_coding | 10 |
| ENSFALG00000010446 | ACAN | protein_coding | 10 |
| ENSFALG00000010524 | TICRR | protein_coding | 10 |
| ENSFALG00000010540 | WDR93 | protein_coding | 10 |
| ENSFALG00000005800 | ARNT2 | protein_coding | 10 |
| ENSFALG00000005835 | CFAP161 | protein_coding | 10 |
| ENSFALG00000011147 | BNC1 | protein_coding | 10 |
| ENSFALG00000002524 | ADAT1 | protein_coding | 11 |
| ENSFALG00000007914 | FTO | protein_coding | 11 |
| ENSFALG00000007821 | GPATCH1 | protein_coding | 11 |
| ENSFALG00000007857 | MCM2 | protein_coding | 12 |
| ENSFALG00000012601 | KLF15 | protein_coding | 12 |
| ENSFALG00000012603 | CFAP100 | protein_coding | 12 |
| ENSFALG00000012626 |  | protein_coding | 12 |
| ENSFALG00000012642 |  | protein_coding | 12 |
| ENSFALG00000012708 |  | protein_coding | 12 |
| ENSFALG00000012724 |  | protein_coding | 12 |
| ENSFALG00000013026 |  | protein_coding | 12 |
| ENSFALG00000014492 | CCDC71 | protein_coding | 12 |
| ENSFALG00000013687 |  | protein_coding | 12 |
| ENSFALG00000007957 | TXNDC15 | protein_coding | 13 |
| ENSFALG00000001023 | FSTL4 | protein_coding | 13 |
| ENSFALG00000001039 |  | protein_coding | 13 |
| ENSFALG00000001103 | P4HA2 | protein_coding | 13 |
| ENSFALG00000003392 | KCNMB1 | protein_coding | 13 |
| ENSFALG00000004993 | CRAMP1 | protein_coding | 14 |
| ENSFALG00000004934 | IFT140 | protein_coding | 14 |
| ENSFALG00000004864 |  | protein_coding | 14 |
| ENSFALG00000014244 |  | protein_coding | 14 |
| ENSFALG00000013840 | AP5Z1 | protein_coding | 14 |
| ENSFALG00000013844 |  | protein_coding | 14 |
| ENSFALG00000013859 | SDK1 | protein_coding | 14 |
| ENSFALG00000014021 | DNAH3 | protein_coding | 14 |
| ENSFALG00000014038 | CCP110 | protein_coding | 14 |
| ENSFALG00000014058 | MARF1 | protein_coding | 14 |
| ENSFALG00000014068 |  | protein_coding | 14 |
| ENSFALG00000014071 |  | protein_coding | 14 |
| ENSFALG00000009070 | KNTC1 | protein_coding | 15 |
| ENSFALG00000008244 | FBRSL1 | protein_coding | 15 |
| ENSFALG00000007995 | CDC45 | protein_coding | 15 |
| ENSFALG00000007409 |  | protein_coding | 15 |
| ENSFALG00000006803 | MORC2 | protein_coding | 15 |
| ENSFALG00000005132 | EXD3 | protein_coding | 17 |
| ENSFALG00000014511 | TRIM32 | protein_coding | 17 |
| ENSFALG00000002222 | RALGPS1 | protein_coding | 17 |
| ENSFALG00000002225 |  | protein_coding | 17 |
| ENSFALG00000009509 |  | protein_coding | 18 |
| ENSFALG00000001307 | MRPL27 | protein_coding | 18 |
| ENSFALG00000001393 | ANKRD40 | protein_coding | 18 |
| ENSFALG00000002286 |  | protein_coding | 18 |
| ENSFALG00000002409 | CACNG1 | protein_coding | 18 |
| ENSFALG00000011945 | DNAH17 | protein_coding | 18 |
| ENSFALG00000011716 | GPS1 | protein_coding | 18 |
| ENSFALG00000011694 | FASN | protein_coding | 18 |
| ENSFALG00000009743 | LIG3 | protein_coding | 19 |
| ENSFALG00000005626 |  | protein_coding | 19 |
| ENSFALG00000005489 | BRIP1 | protein_coding | 19 |
| ENSFALG00000005200 |  | protein_coding | 19 |
| ENSFALG00000012472 | CEP250 | protein_coding | 20 |
| ENSFALG00000012569 |  | protein_coding | 20 |
| ENSFALG00000012591 |  | protein_coding | 20 |
| ENSFALG00000005773 | RAD21L1 | protein_coding | 20 |
| ENSFALG00000005885 |  | protein_coding | 20 |
| ENSFALG00000012924 |  | protein_coding | 20 |
| ENSFALG00000012895 | NCOA3 | protein_coding | 20 |
| ENSFALG00000003840 |  | protein_coding | 21 |
| ENSFALG00000003676 | EIF4G3 | protein_coding | 21 |
| ENSFALG00000003551 |  | protein_coding | 21 |
| ENSFALG00000007239 | UBE4B | protein_coding | 21 |
| ENSFALG00000006422 | KAT6A | protein_coding | 22 |
| ENSFALG00000009791 | C2CD2L | protein_coding | 24 |
| ENSFALG00000009812 | VPS11 | protein_coding | 24 |
| ENSFALG00000009912 | ARCN1 | protein_coding | 24 |
| ENSFALG00000009959 |  | protein_coding | 24 |
| ENSFALG00000009979 | UBE4A | protein_coding | 24 |
| ENSFALG00000010009 |  | protein_coding | 24 |
| ENSFALG00000000766 | MAGI3 | protein_coding | 26 |
| ENSFALG00000000504 |  | protein_coding | 26 |
| ENSFALG00000003157 |  | protein_coding | 26 |
| ENSFALG00000002495 |  | protein_coding | 26 |
| ENSFALG00000002397 |  | protein_coding | 26 |
| ENSFALG00000014315 | GJC1 | protein_coding | 27 |
| ENSFALG00000014101 |  | protein_coding | 27 |
| ENSFALG00000015289 |  | protein_coding | 27 |
| ENSFALG00000000804 | NBR1 | protein_coding | 27 |
| ENSFALG00000000796 | BRCA1 | protein_coding | 27 |
| ENSFALG00000000389 | TOP2A | protein_coding | 27 |
| ENSFALG00000000467 |  | protein_coding | 27 |
| ENSFALG00000013413 | EPS15L1 | protein_coding | 28 |
| ENSFALG00000012190 | PRRC1 | protein_coding | KE165356.1 |
| ENSFALG00000002791 |  | protein_coding | KE165377.1 |
| ENSFALG00000009297 |  | protein_coding | KE165418.1 |
| ENSFALG00000013251 | RBL2 | protein_coding | KE165366.1 |
| ENSFALG00000001189 |  | protein_coding | KE165444.1 |
| ENSFALG00000011832 |  | protein_coding | KE165373.1 |
| ENSFALG00000013276 | CHD9 | protein_coding | KE165366.1 |
| ENSFALG00000002800 |  | protein_coding | KE165377.1 |
| ENSFALG00000009773 | MALT1 | protein_coding | AGTO02007302.1 |
| ENSFALG00000004561 |  | protein_coding | AGTO02006776.1 |
| ENSFALG00000008996 |  | protein_coding | Z |
| ENSFALG00000007391 | DMXL1 | protein_coding | KE165325.1 |
| ENSFALG00000010499 |  | protein_coding | KE165357.1 |
| ENSFALG00000000047 | APC | protein_coding | KE165353.1 |
| ENSFALG00000013069 | FHDC1 | protein_coding | KE165308.1 |
| ENSFALG00000003397 | KDM3B | protein_coding | KE165350.1 |
| ENSFALG00000007573 | SNCAIP | protein_coding | KE165343.1 |
| ENSFALG00000013075 | MND1 | protein_coding | KE165308.1 |
| ENSFALG00000001153 | MEGF10 | protein_coding | KE165345.1 |
| ENSFALG00000007585 |  | protein_coding | KE165343.1 |
| ENSFALG00000012728 | ATRN | protein_coding | KE165344.1 |
| ENSFALG00000013125 |  | protein_coding | KE165308.1 |
| ENSFALG00000001044 |  | protein_coding | KE165324.1 |
| ENSFALG00000008928 | KATNAL2 | protein_coding | Z |
| ENSFALG00000006476 | FREM1 | protein_coding | KE165323.1 |
| ENSFALG00000006534 | PSIP1 | protein_coding | KE165323.1 |
| ENSFALG00000006548 | CCDC171 | protein_coding | KE165323.1 |
| ENSFALG00000009881 | KIF24 | protein_coding | Z |
| ENSFALG00000009799 |  | protein_coding | Z |
| ENSFALG00000002166 |  | protein_coding | Z |
| ENSFALG00000002091 | SPEF2 | protein_coding | Z |
| ENSFALG00000002085 | IL7R | protein_coding | Z |
| ENSFALG00000002083 | CAPSL | protein_coding | Z |
| ENSFALG00000000209 | SMARCA1 | protein_coding | 4A |
| ENSFALG00000002055 |  | protein_coding | Z |
| ENSFALG00000000199 |  | protein_coding | 4A |
| ENSFALG00000000187 | COL4A6 | protein_coding | 4A |
| ENSFALG00000002044 | EGFLAM | protein_coding | Z |
| ENSFALG00000000173 | ACSL4 | protein_coding | 4A |
| ENSFALG00000002041 | LIFR | protein_coding | Z |
| ENSFALG00000000168 | TMEM164 | protein_coding | 4A |
| ENSFALG00000000157 |  | protein_coding | 4A |
| ENSFALG00000002027 | FYB1 | protein_coding | Z |
| ENSFALG00000002020 | DAB2 | protein_coding | Z |
| ENSFALG00000012116 | MAMLD1 | protein_coding | 4A |
| ENSFALG00000001961 | C6 | protein_coding | Z |
| ENSFALG00000011136 | NIM1K | protein_coding | Z |
| ENSFALG00000011164 | C5orf34 | protein_coding | Z |
| ENSFALG00000011192 | NNT | protein_coding | Z |
| ENSFALG00000010749 | ITGA2 | protein_coding | Z |
| ENSFALG00000010796 |  | protein_coding | Z |
| ENSFALG00000010919 | IL31RA | protein_coding | Z |
| ENSFALG00000010893 | ATRX | protein_coding | 4A |
| ENSFALG00000011027 | DEPDC1B | protein_coding | Z |
| ENSFALG00000009671 | MOV10L1 | protein_coding | 1A |
| ENSFALG00000011073 | KIF2A | protein_coding | Z |
| ENSFALG00000011086 | IPO11 | protein_coding | Z |
| ENSFALG00000009908 |  | protein_coding | 1A |
| ENSFALG00000009863 | CENPH | protein_coding | Z |
| ENSFALG00000014553 |  | protein_coding | Z |
| ENSFALG00000010196 | ARHGEF28 | protein_coding | Z |
| ENSFALG00000010206 | UTP15 | protein_coding | Z |
| ENSFALG00000010312 |  | protein_coding | Z |
| ENSFALG00000010416 | SMARCA2 | protein_coding | Z |
| ENSFALG00000010509 | PUM3 | protein_coding | Z |
| ENSFALG00000010531 |  | protein_coding | Z |
| ENSFALG00000010553 |  | protein_coding | Z |
| ENSFALG00000010582 | RIC1 | protein_coding | Z |
| ENSFALG00000010612 | KIAA2026 | protein_coding | Z |
| ENSFALG00000003044 | BNC2 | protein_coding | Z |
| ENSFALG00000003055 | ADAMTSL1 | protein_coding | Z |
| ENSFALG00000003066 | PLIN2 | protein_coding | Z |
| ENSFALG00000003075 | DENND4C | protein_coding | Z |
| ENSFALG00000003119 | FOCAD | protein_coding | Z |
| ENSFALG00000003147 |  | protein_coding | Z |
| ENSFALG00000003176 |  | protein_coding | Z |
| ENSFALG00000003541 | TUT7 | protein_coding | Z |
| ENSFALG00000003400 |  | protein_coding | Z |
| ENSFALG00000004759 | SYK | protein_coding | Z |
| ENSFALG00000005018 |  | protein_coding | Z |
| ENSFALG00000010495 | OTOGL | protein_coding | 1A |
| ENSFALG00000005146 | HOOK3 | protein_coding | Z |
| ENSFALG00000001025 |  | protein_coding | Z |
| ENSFALG00000000968 |  | protein_coding | Z |
| ENSFALG00000001173 | TDRD7 | protein_coding | Z |
| ENSFALG00000001223 | DNAJA1 | protein_coding | Z |
| ENSFALG00000007333 | TEK | protein_coding | Z |
| ENSFALG00000015114 |  | protein_coding | Z |
| ENSFALG00000000186 |  | protein_coding | Z |
| ENSFALG00000000166 | SHOC1 | protein_coding | Z |
| ENSFALG00000000118 | SLC46A2 | protein_coding | Z |
| ENSFALG00000000116 |  | protein_coding | Z |
| ENSFALG00000002623 |  | protein_coding | Z |
| ENSFALG00000002553 | ZNF608 | protein_coding | Z |
| ENSFALG00000004755 | PTAR1 | protein_coding | Z |
| ENSFALG00000012713 | TRPM6 | protein_coding | Z |
| ENSFALG00000002900 | FSD1L | protein_coding | Z |
| ENSFALG00000000833 |  | protein_coding | Z |
| ENSFALG00000009043 | ADGRV1 | protein_coding | Z |
| ENSFALG00000009074 | KIAA0825 | protein_coding | Z |
| ENSFALG00000009108 |  | protein_coding | Z |
| ENSFALG00000012891 | VCAN | protein_coding | Z |
| ENSFALG00000000544 | SLC13A4 | protein_coding | 1A |
| ENSFALG00000000511 | CREB3L2 | protein_coding | 1A |
| ENSFALG00000002808 | SMC1B | protein_coding | 1A |
| ENSFALG00000001086 | PLCZ1 | protein_coding | 1A |
| ENSFALG00000000125 | NA | NA | NA |
| ENSFALG00000000138 | NA | NA | NA |
| ENSFALG00000000534 | NA | NA | NA |
| ENSFALG00000000783 | NA | NA | NA |
| ENSFALG00000001041 | NA | NA | NA |
| ENSFALG00000002263 | NA | NA | NA |
| ENSFALG00000002584 | NA | NA | NA |
| ENSFALG00000002858 | NA | NA | NA |
| ENSFALG00000003146 | NA | NA | NA |
| ENSFALG00000003236 | NA | NA | NA |
| ENSFALG00000003286 | NA | NA | NA |
| ENSFALG00000003956 | NA | NA | NA |
| ENSFALG00000004569 | NA | NA | NA |
| ENSFALG00000004815 | NA | NA | NA |
| ENSFALG00000005158 | NA | NA | NA |
| ENSFALG00000005411 | NA | NA | NA |
| ENSFALG00000005722 | NA | NA | NA |
| ENSFALG00000006158 | NA | NA | NA |
| ENSFALG00000007897 | NA | NA | NA |
| ENSFALG00000008034 | NA | NA | NA |
| ENSFALG00000009077 | NA | NA | NA |
| ENSFALG00000009208 | NA | NA | NA |
| ENSFALG00000009891 | NA | NA | NA |
| ENSFALG00000009894 | NA | NA | NA |
| ENSFALG00000010067 | NA | NA | NA |
| ENSFALG00000010084 | NA | NA | NA |
| ENSFALG00000010186 | NA | NA | NA |
| ENSFALG00000010474 | NA | NA | NA |
| ENSFALG00000010516 | NA | NA | NA |
| ENSFALG00000010613 | NA | NA | NA |
| ENSFALG00000011116 | NA | NA | NA |
| ENSFALG00000012506 | NA | NA | NA |
| ENSFALG00000012531 | NA | NA | NA |
| ENSFALG00000013613 | NA | NA | NA |
| ENSFALG00000013903 | NA | NA | NA |
| ENSFALG00000014353 | NA | NA | NA |
| ENSFALG00000015115 | NA | NA | NA |
| ENSFALG00000015151 | NA | NA | NA |
| ENSFALG00000015261 | NA | NA | NA |
